# Structural Insights into Broad-Range Polyphosphate Kinase 2-II Enzymes Applicable for Pyrimidine Nucleoside Diphosphate Synthesis

**DOI:** 10.1101/2024.11.26.625422

**Authors:** Marco Kuge, Michael Keppler, Florian Friedrich, Raspudin Saleem-Batcha, Juliana Winter, Isabel Prucker, Philipp Germer, Stefan Gerhardt, Oliver Einsle, Manfred Jung, Henning J. Jessen, Jennifer N. Andexer

## Abstract

Polyphosphate kinases (PPK) play crucial roles in various biological processes, including energy storage and stress responses, through their interaction with inorganic polyphosphate (polyP) and the intracellular nucleotide pool. Members of the PPK family 2 (PPK2s) catalyse polyP-consuming phosphorylation of nucleotides. In this study, we characterised two PPK2 enzymes from *Bacillus cereus* (*Bc*PPK2) and *Lysinibacillus fusiformis* (*Lf*PPK2) to investigate their substrate specificity and potential for selective nucleotide synthesis. Both enzymes exhibited a broad substrate scope, selectively converting over 85% of pyrimidine nucleoside monophosphates (NMPs) to nucleoside diphosphates (NDPs), while nucleoside triphosphate (NTP) formation was observed only with purine NMPs. Preparative enzymatic synthesis of cytidine diphosphate (CDP) was applied to achieve an yield of 49%. Finally, structural analysis of five crystal structures of *Bc*PPK2 and *Lf*PPK2 provided insights into their active sites and substrate interactions. This study highlights PPK2-II enzymes as promising biocatalysts for the efficient and selective synthesis of pyrimidine NDPs.

## Introduction

Inorganic polyphosphate (polyP) is an omnipresent molecule conserved in all domains of life. The polymer, containing up to hundreds of phosphate groups, is linked by high energy phosphoanhydride bonds.^[1]^ These energy-rich bonds render polyP a common polymer for energy and phosphate storage in various organisms.^[2]^ Additionally, polyP is associoated with virulence and biofilm production of different bacteria and archaea as well as stress response mechanisms.^[3–5]^ In 1956, Kornberg *et al*. first described the nucleoside triphosphate (NTP) dependent synthesis of polyP catalysed by polyP kinase (PPK) in *Escherichia coli*.^[6]^ When a new enzyme family of polyP-dependent nucleotide kinases was described in *Pseudomonas aeruginosa*, the *E. coli* enzyme was later annotaded as family 1 PPK (PPK1), while the novel enzyme family was classified as family 2 PPK (PPK2).^[7]^ PPK1 and PPK2 are structurally different, the latter have so far not been described in mammalian genomes.^[8]^

Based on sequence characteristics and substrate preferences PPK2s are further subdivided into three classes: PPK2-I, PPK2-II and PPK2-III have been described to catalyse the phosphorylation of nucleoside diphosphates (NDPs), nucleoside monophosphates (NMPs), or both, respectively (Figure 1).^[9]^ Our studies suggest that these substrate range definitions are a result of kinetic preferences, as all classes demonstrated acceptance of all phosphorylation states under extended reaction times.^[10,11]^ Nevertheless, these preferences offer unique possibilities to control the outcome of nucleotide phosphorylation reactions. PPK2 enzymes can be further classified into one-or two-domain enzymes, with the latter consisting of two fused PPK2 domains, probably resulting from a gene duplication event. The classes PPK2-I and -III consist of one-domain enzymes, the PPK2-II class includes both one-and two-domain proteins.^[9]^ Expanding our understanding about the substrate scope and selectivity of different representatives of PPK2 enzymes has the potential to unlock access to a variety of nucleotide building blocks in their desired phosphorylation state. As one example, Eltouhky *et al*. used either a PPK2-II or PPK2-III enzyme to selectively synthesise the NDP or NTP, respectively, of acadesine, a nucleoside analogue drug in the treatment of leukemia.^[12]^

**Figure 1.**
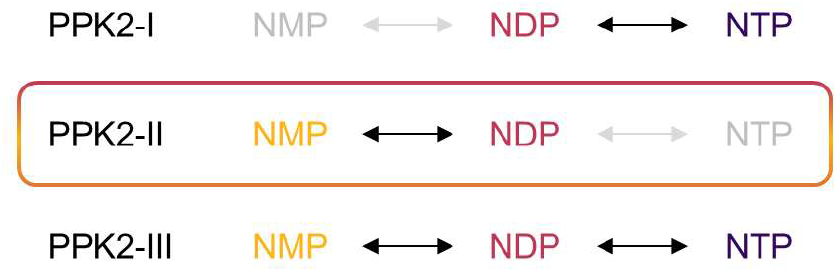
PPK2 enzymes are divided into three subclasses. Each subclass (PPK2-I, -II, -III) shows kinetic preferences for the polyP-dependent phosphorylation of nucleotides in different phosphorylation states. This work focusses on PPK2-II (coloured box).

In the past, crystal structures of several bacterial PPK2s have been solved and biochemical characterisations of representatives of PPK2-I to –III have been reported.^[13–15]^ Most of the studies demonstrated the enzymes’ ability to accept purine nucleotides, such as adenosine mono-or diphosphate (AMP/ADP). In general, pyrimidine nucleotides, including cytidine mono- or diphosphate (CMP/CDP), show lower substrate acceptance and turnover rates with most PPK2 enzymes.^[16,17]^ Examples for more promiscuous PPK2s are PPK2c from *Ralstonia eutropha* and PPK3 from *Ruegeria pomeroyi*. Both enzymes accept purine and pyrimidine diphosphates.^[18,19]^ Sequence characteristics identify both as PPK2-I enzymes. Additionally, a PPK2-III from *Meiothermus ruber* accepts mono- and diphosphorylated purine-and pyrimidine-derived nucleotides.^[9]^

Given their ability to catalyse NTP formation, most research on PPK2 enzymes focused on PPK2-I and PPK2-III and their utilisation in ATP regeneration systems.^[20–22]^ In recent years, non-canonical purine and pyrimidine nucleotide analogues are gaining significance in synthetic biology, with wide-spread applications in drug development. For some application, pyrimidine nucleotides are especially relevant, e.g. in the biosynthesis of thiamine (vitamin B_1_) or in the generation of sugars including sialic acids.^[23–25]^ Pyrimidine nucleotide analogues are established treatments for various viral diseases, such as AIDS or infectious mononucleosis, demonstrating a variety of desirable pharmacological activities, including antibacterial, antifungal and anticancer effects.^[26,27]^

PPK2 enzymes are a valuable tool for nucleotide synthesis and further research in this topic contributes towards their application. With only one solved crystal structure of a two-domain PPK2-II enzyme, *Pa*PPK2-II (from *Pseudomonas aeruginosa*, PDB ID 3CZP) much remains unknown about this enzyme class and differences to PPK2-I and -III.^[16]^ Additionally, the only PPK2-II described to accept pyrimidine-derived NMPs was isolated from *Acinetobacter johnsonii* (*Aj*PPK2-II). However, its activity on pyrimidine-derived nucleotides was three orders of magnitude lower compared to purine nucleotides.^[29]^

This study aims to characterise novel PPK2-II enzymes to facilitate selective synthesis of NDPs. Direct NDP synthesis has been described as difficult by classical chemical approaches.^[30,31]^ We identified two one-domain PPK2-II enzymes from *Bacillus cereus* (*Bc*PPK2) and *Lysinibacillus fusiformis* (*Lf*PPK2) as suitable candidates and investigated them in more detail (Table 1). While the latter has not been described in literature before, *Bc*PPK2-II has been linked to biofilm formation and motility in knockout studies.^[32]^

**Table 1.** PPK2-II enzymes characterised in this work.

| Enzyme | Organism | Sequence identity to AjPPK2 [%]<sup>[a]</sup> | UniProt ID |
| --- | --- | --- | --- |
| AjPPK2 | *Acinetobacter johnsonii* | - | Q83XD3 |
| BcPPK2 | *Bacillus cereus* | 45.8 | Q81F46 |
| LfPPK2 | *Lysinibacillus fusiformis* | 47.0 | A0A1E4R1F9 |

[a] Alignment performed with Clustal Omega (Figure S9).<sup>[28]</sup>

## Results and Discussion

### *Bc*PPK2 and *Lf*PPK2 Show Broad Substrate Ranges

Following the successful production and purification of *Bc*PPK2 and *Lf*PPK2 (Figure S1), an activity assay using AMP as the substrate was conducted.

Reaction products were analysed *via* capillary electrophoresis coupled with electron spray ionisation mass spectroscopy (CE-ESI-MS).^[33,34]^ Reaction conditions were adapted from previous work with PPK2-I from *Sinorhizobium meliloti*.^[11]^ Starting from AMP, both ADP and ATP were detected in both experiments, with ADP as the main product (Figure 2). After the confirmation of catalytic activity, we conducted a substrate screening based on HPLC-UV/Vis detection to test both biocatalysts for their acceptance towards different NMPs. This substrate panel included purine, pyrimidine, and 2’-deoxy NMPs as potential phosphate acceptors (Figure 3 and S5). The aim was to characterise the substrate promiscuity of the two PPK2-II enzymes as well as their selectivity in producing nucleotides in different phosphorylation states.

**Figure 2.**
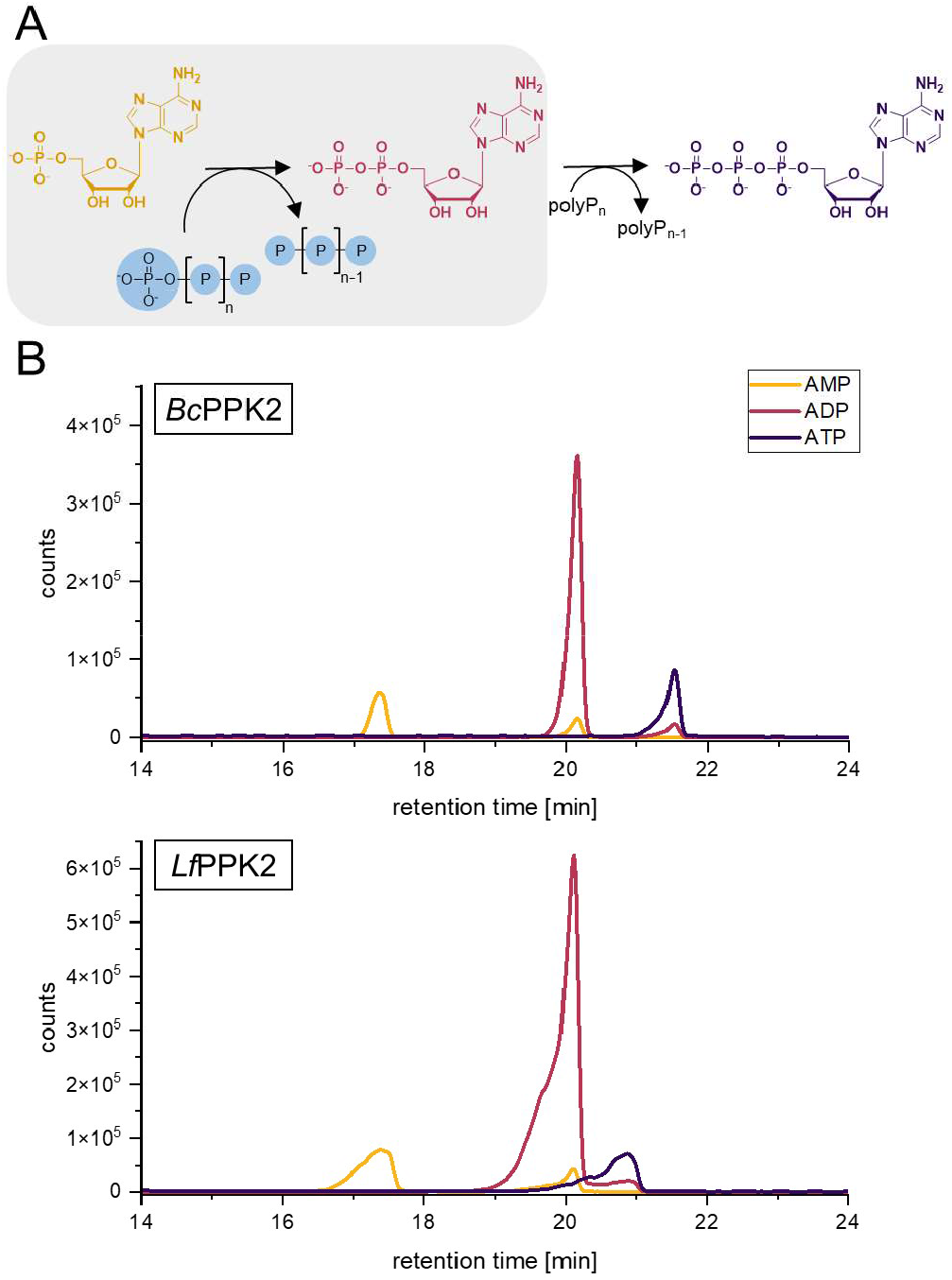
Reaction scheme of polyP-dependent phosphorylation of AMP (A). Extracted ion electropherogram of CE-ESI-MS measurements of enzyme activity assay with *Bc*PPK2 and *Lf*PPK2 (B). Masses corresponding to AMP (yellow), ADP (red) and ATP (purple) can be observed with ADP being the main reaction product. Peak assignment was done *via* MS/MS transitions (Table S3). Additional peaks of AMP and ADP below main peaks are results of neutral loss of phosphate during ionisation.

**Figure 3.**
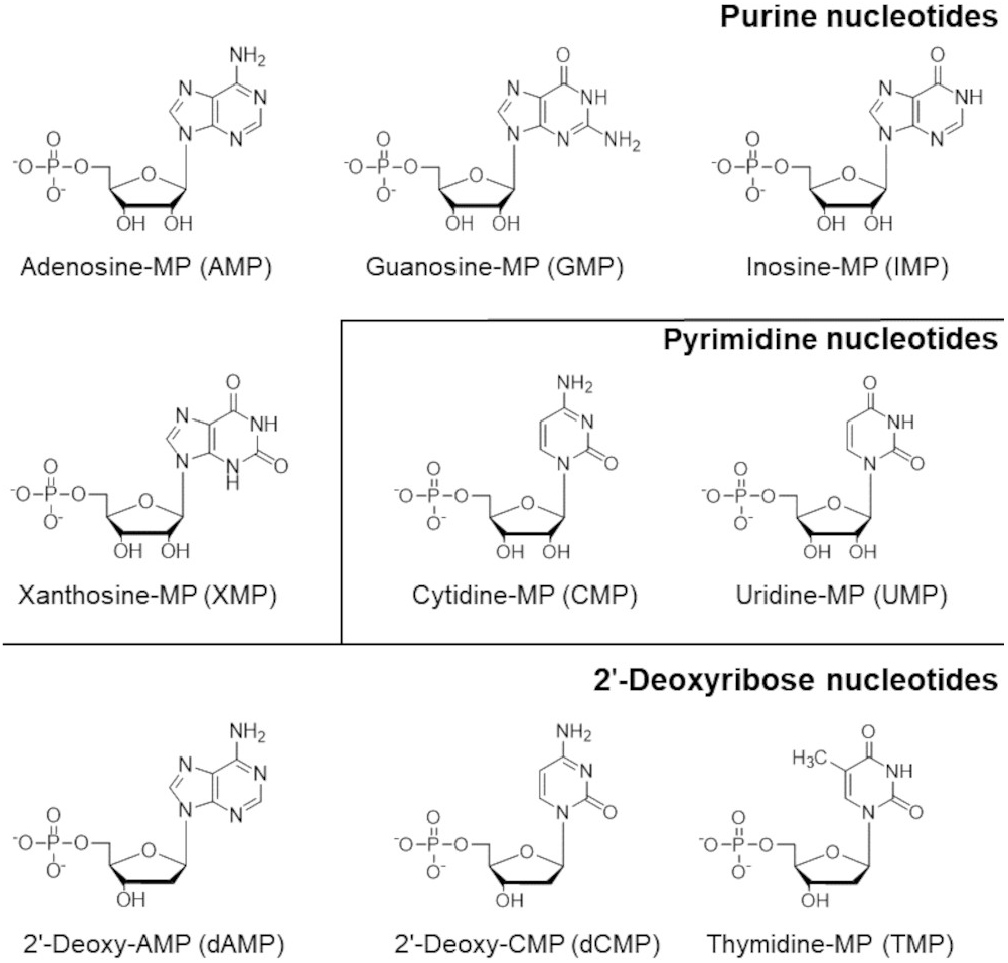
Panel of NMPs used for substrate screening.

All tested (d)NMPs were accepted by both enzymes revealing an exceptionally broad substrate range with a clear tendency to form the corresponding NDP as the main product (Figure 4). Interestingly, substrate-specific phosphorylation was observed. While NTP formation was observed for purine NMPs, exclusive monophosphorylation of pyrimidine and 2′-deoxy NMPs to the corresponding NDPs was detected. ATP formation up to 28% for *Bc*PPK2 and 20% for *Lf*PPK2 was measured. GTP formation was lower, with about 10% for both enzymes. Both enzymes catalysed the phosphorylation of the pyrimidine NMPs CMP and UMP to the corresponding NDPs.In our setup conversion of 85% of CMP and UMP as well as 60% of TMP to the corresponding NDPs were observed. The lowest conversion was observed for dAMP, with 57% NDP formation for *Bc*PPK2 and 17% for *Lf*PPK2. Overall, both *Bc*PPK2 and *Lf*PPK2 yield similar amounts of synthesised NDPs (and NTPs).

**Figure 4.**
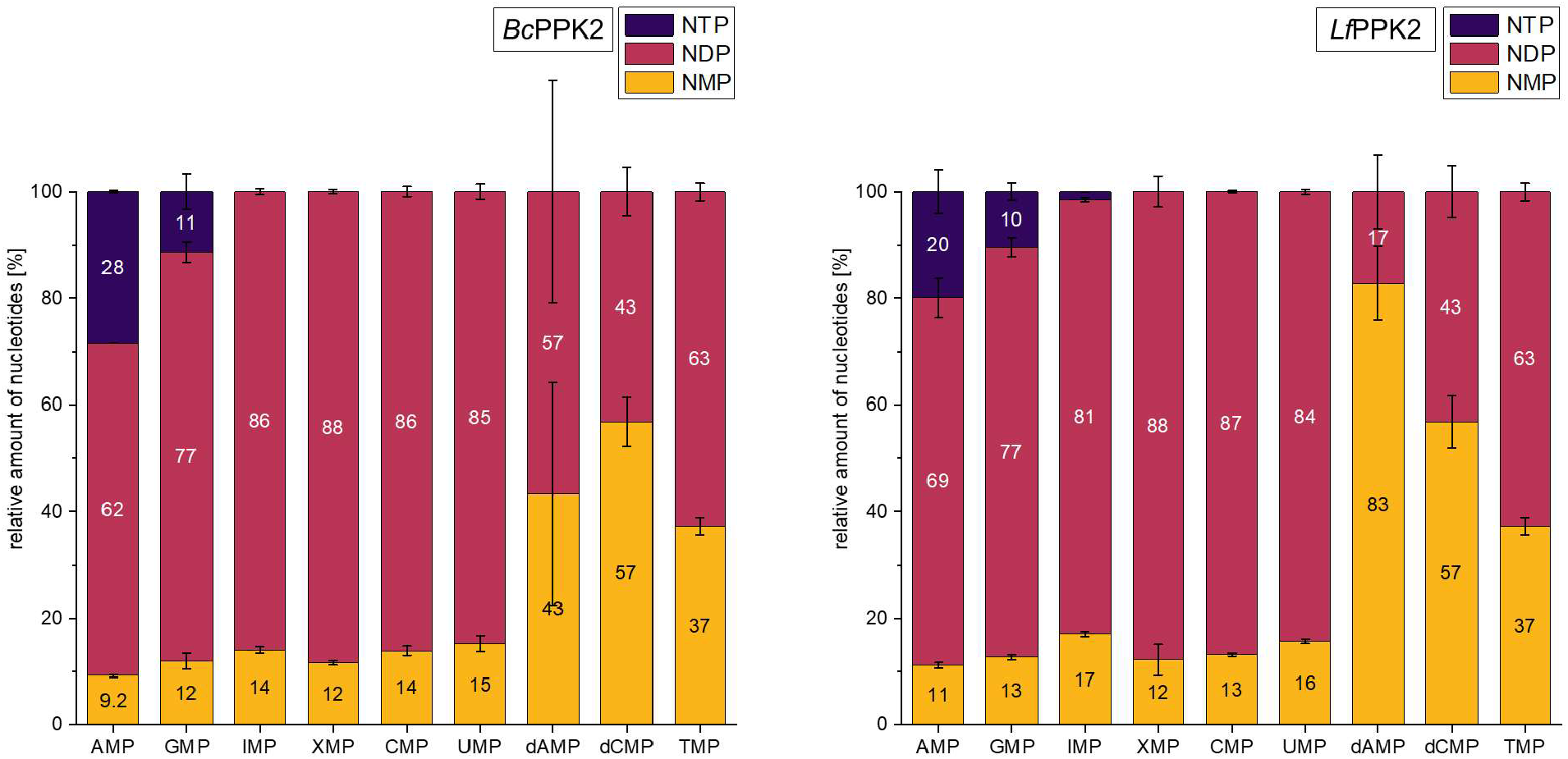
Substrate specificity of *Bc*PPK2-II and *Lf*PPK2-II. Enzyme assays comprised 1 µM PPK2 enzyme, 1 mM NMP, 10 mM MgCl_2_, 10 mM polyP (calculated as single phosphate units) and were incubated at 37 °C for 20 h. After quenching with perchloric acid (2.5% final concentration), samples were subjected to HPLC analysis.

### *Lf*PPK2-II Converts Pyrimidine Nucleotide CMP Faster than *Aj*PPK2

For *Aj*PPK2-II, Shiba *et al*. reported a relative activity of 0.09% on CMP compared to its preferred substrate AMP.^[29]^ We conducted a time course experiment with *Aj*PPK2 and *Lf*PPK2 as biocatalysts for the phosphorylation of CMP. In our reaction setup both enzymes readily used CMP as phosphate acceptor to form CDP. The outcome after 20 h incubation is similar for both proteins, with a relative conversion of 85% for *Lf*PPK2 and 83% for *Aj*PPK2 (Figure 5). Again, triphosphate formation was not observed. For confirmation, experiments were repeated with CDP as starting substrate, resulting in the same ratio of CMP to CDP with no observed CTP formation (Figure S2). Nevertheless, the proteins differ in their velocity. *Lf*PPK2 converted over 50% of CMP to CDP after 120 minutes, while *Aj*PPK2 shows 30% conversion after 120 minutes and around 40% after 240 minutes. For the preferred substrate AMP, both enzymes show a substantially higher velocity (85% of conversion after 10 minutes already, Figure S3). ATP formation was first observed after 120 min for both enzymes. After 20 h *Lf*PPK2 and *Aj*PPK2 phosphorylated 20% and 38% of AMP to ATP, respectively.

**Figure 5.**
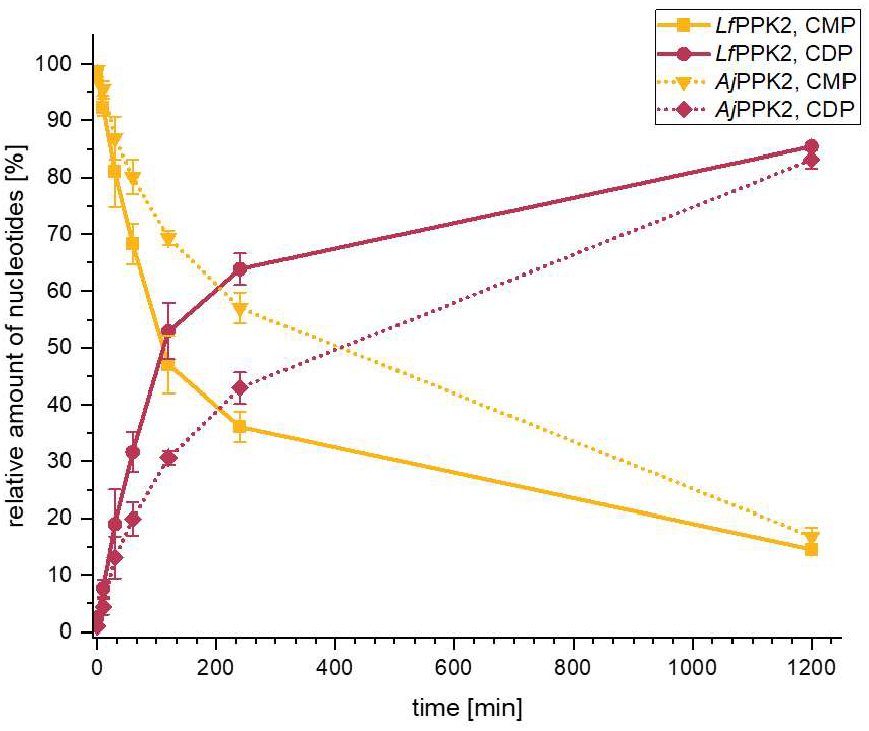
Time course of CDP formation catalysed by *Lf*PPK2 and *Aj*PPK2. Assays comprised 1 mM CMP, 1 µM PPK2 enzyme, 10 mM polyP, 10 mM MgCl_2_. Samples were taken after 0, 10, 30, 60, 120, 240 and 1200 minutes and subjected to HPLC analysis.

### PPK2-II Enzymes Enable Access to Pyrimidine NDPs *via* Ion Exchange Chromatography

To test the suitability of PPK2-II enzymes for the polyP-dependent preparative synthesis of pyrimidine NDPs, CMP was chosen as model substrate and *Lf*PPK2 as biocatalyst. The reaction was set up in 5 mL scale with 10 mM CMP final concentration. The assay was conducted for 20 h, filtered and applied onto a HiPrep Q FF 20 mL anion exchange column. Nucleotides were eluted with a gradient of NaClO_4_, pH 4.2 (Figure 6).

**Figure 6.**
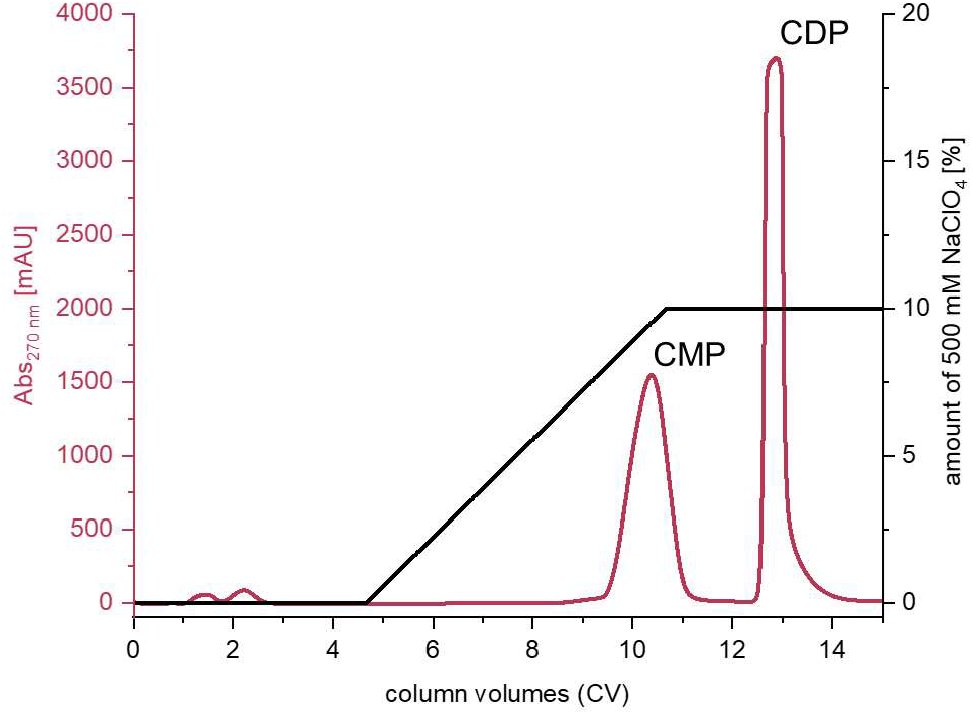
Ion exchange chromatogram obtained during preparation of CDP. The assay comprised 5 µM *Lf*PPK2, 10 mM CMP, 10 mM polyP (calculated as single phosphate units), 10 mM MgCl_2_ in 50 mM TRIS, pH 8.0.

The pooled fractions containing the desired CDP were purified as previously described for purine NTPs *via* precipitation in acetone.^[35]^ The conversion was 54% (determined *via* peak integration), yielding 8.5 mg of CDP as the sodium salt after purification. The identity of CDP was confirmed in ^1^H and ^31^P nuclear magnectic resonance (NMR) measurements as well as HPLC analysis compared to an authentic CDP standard (Figure S6-S8). The conversion of 54% differs from the previous small-scale substrate screening experiment (87% conversion, Figure 5). As product inhibition is not uncommon in upscaled reactions, this might be a contributing factor and can be addressed by reaction engineering.^[36,37]^

### Crystal Structures of *Bc*PPK2 and *Lf*PPK2

To gain further understanding of the exceptional broad substrate scope of *Bc*PPK2 and *Lf*PPK2, we conducted several crystallisation experiments with both proteins. Utilising the sitting drop vapor diffusing method, co-crystallisation and soaking approaches we report a total of five crystal structures (Table 2, S1, S2). These represent the first examples of substrate-bound PPK2-II enzyme structures solved, as well as the first PPK2 structures in complex with 2′-deoxy or pyrimidine nucleotides of any subclass.

**Table 2.** Summary of crystal structures solved in this study.

| Enzyme | Substrate | Resolution [Å] | Data Completeness [%] | $R_{free}$ | PDB ID |
| --- | --- | --- | --- | --- | --- |
| *BcPPK2* | - | 2.55 | 99.7 | 0.25 | 9GIA |
| *LfPPK2* | - | 2.20 | 98.7 | 0.25 | 9H8J |
| *LfPPK2* | AMP | 2.10 | 99.5 | 0.22 | 9H8K |
| *LfPPK2* | ADP | 2.88 | 98.6 | 0.23 | 9GP9 |
| *LfPPK2* | TMP | 2.10 | 96.6 | 0.21 | 9H8L |

The overall structure of both enzymes is consistent with published structures of all three PPK2 classes. The apo structures have the typical three-layered α/β/α-sandwich, where five β-sheets are flanked by three α-helices on the one side and four on the other (Figure 7A, 5B and S10). The structures show the characteristic lid domain fully resolved, which is noteworthy as PPK2 lid domains are inherently flexible and therefore often lack electron density definition.^[16,38]^ The P-loop kinase conserved motifs Walker A (GxDxxGK) and Walker B (DRS) as well as the PPK2-II signature Gly^[13,15]^ (G114 in *Bc*PPK2, G112 in *Lf*PPK2) are fully resolved for both enzyme structures (Figure 7). PPK2 enzymes have an overall positively charged polyP tunnel with a large number of arginine and lysine residues. This characteristic is also observed in both proteins.

**Figure 7.**
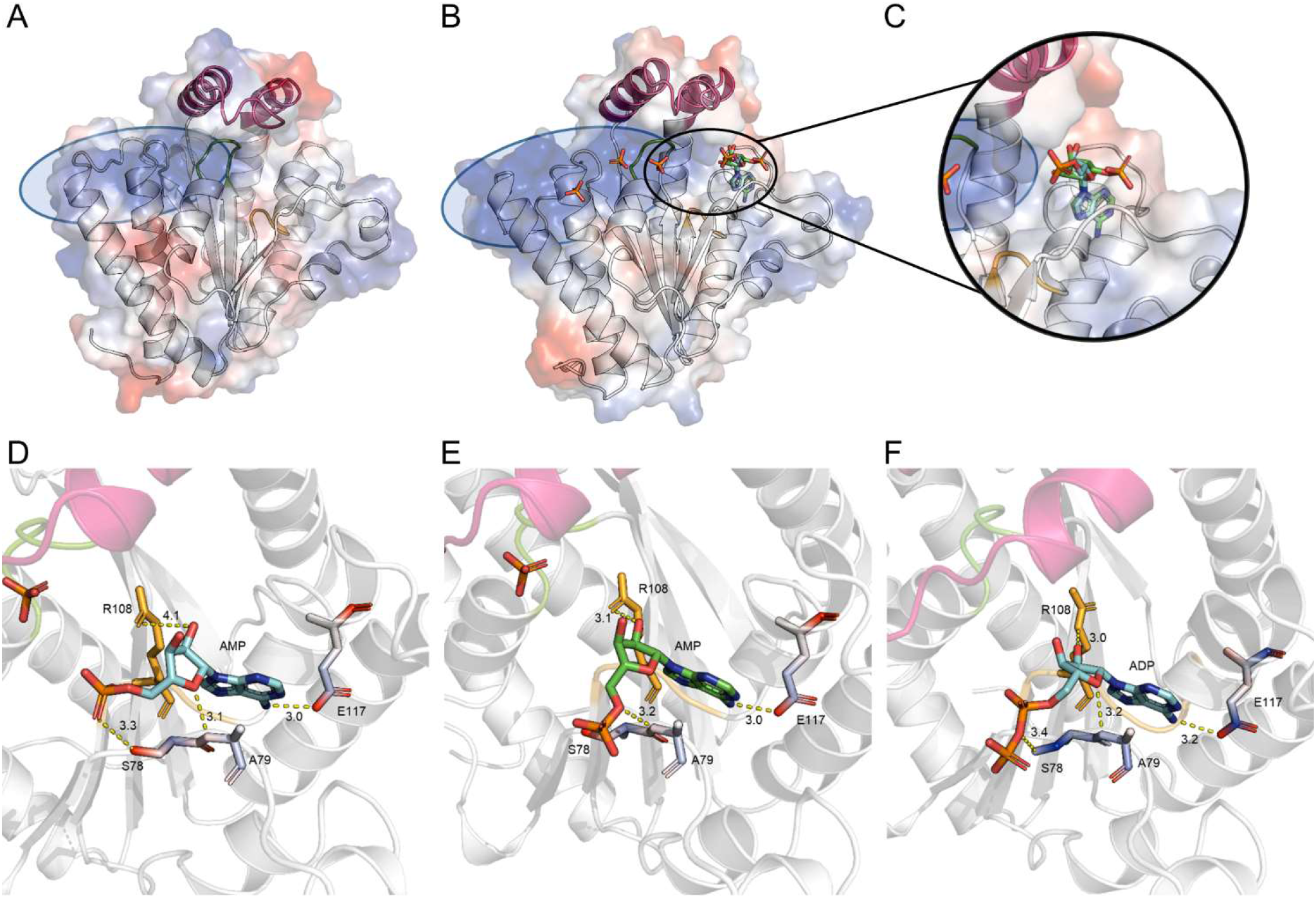
Structure representation of *Bc*PPK2 (A) and *Lf*PPK2 bound to AMP in two conformations (depicted in turquoise and green, B) with closeup view of the substrate AMP in two different conformations (C). Electrostatic surface representation indicating positively charged residues in blue and negatively charged residues in red. Tertiary structure elements are depicted in an overlaid cartoon representation. The conserved motifs are coloured as follows: Walker A, green. Walker B, orange. Lid domain, violet. The overall positively charged polyP tunnel is framed in light blue.Detailed view on substrate binding of *Lf*PPK2 to AMP with its phosphate chain facing towards the polyP tunnel (D), away from it (E) and to ADP (F). Hydrogen bonds are coloured in yellow. A sequence alignment of different PPK2 enzymes and their characteristic motifs can be found in the supporting information (Figure S9).

In the substrate-bound structures with AMP and TMP, a phosphate is coordinated by residues of the Walker A motif (D52-K56), as well as by the adjacent G57 and R168 of the lid domain (Figure 8A and S11). Interestingly, the crystal structure of *Lf*PPK2 bound to AMP shows electron density for two conformations of the nucleotide, one with the phosphate facing towards the polyP tunnel and one facing away from it (Figure 7C). This suggests a high degree of flexibility of the phosphate side chain. The AMP conformer facing towards the polyP tunnel shows four interactions with the enzyme (Figure 7D).

**Figure 8.**
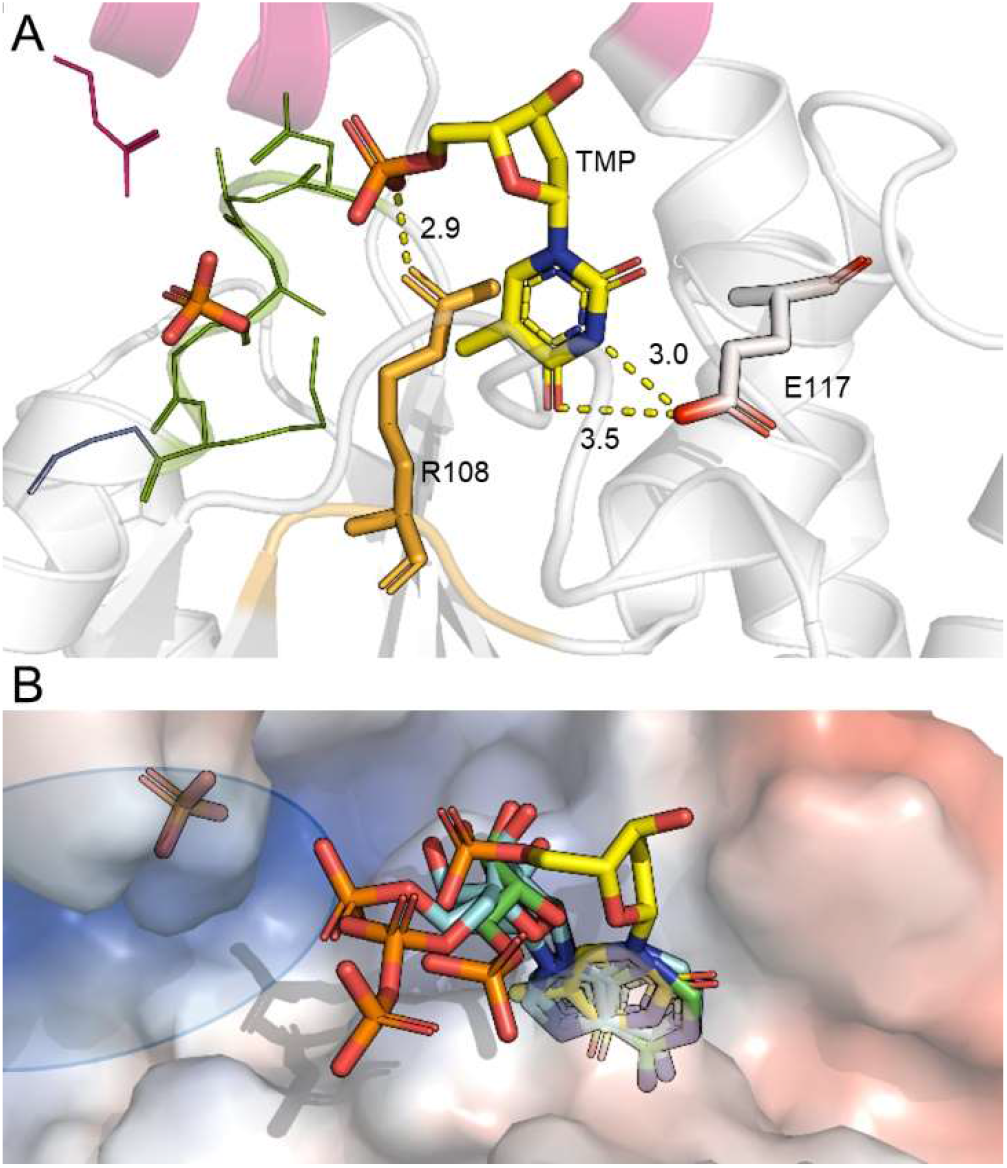
Detailed view on substrate binding of *Lf*PPK2 with TMP (depicted in yellow) (A). Overlay of the substrates AMP (2 confomers in tuquiose and green), ADP (turquoise) and TMP (yellow) (B). The polyP tunnel of *Lf*PPK2 is framed in blue.

The amino group in the C6-position of the adenine nucleobase forms a hydrogen bond with the carboxylate group of E117, at a distance of 3.0 Å. This interaction can be found in all substrate-bound structures of this study. The ribofuranose oxygen interacts with the protein backbone between S78 and A79 (3.1 Å) as well as with R108 (4.1 Å) *via* the hydroxyl group in C2′-position. R108 is part of the Walker B motif and was previously shown to be essential for the catalytic activity of PPK2 enzymes.^[13]^ The phosphate group shows ionic interactions with the side chain of S78 (3.3 Å). The second AMP conformer with its phosphate chain facing away from the polyP tunnel interacts similarly with R108 and E117. In this state, the ribofuranose oxygen is no longer in contact with the protein backbone and the phosphoryl oxygen close to the C5′-position takes its place (3.2 Å). The ribose is slightly tilted so that the C2′ hydroxyl group is now closer to R108 (3.1 Å instead of 4.1 Å). Even though the mechanism of PPK2-II-catalysed phosphorylation is not yet fully understood, previous studies imply that proximity of the nucleotide’s phosphate chain towards the phosphate donor polyP in the polyP tunnel is crucial for the phosphate transfer.^[13,14]^ Therefore, we suggest that the first AMP conformer is meeting this criterion more effectively than the second one. It should be noted that other factors, such as the dependency of PPK2 enzymes on divalent metal ions and the high flexibility of the lid domain, must be taken into account. The observation that the phosphate-chain bearing end seems to retain more flexibility than the nucleobase moiety might be an important factor towards the substrate specifity of PPK2-II enzymes. In the *Lf*PPK2 structure bound to ADP, the nucleotide is interacting with the same residues as AMP (S78, A79, R108, E117). The oxygen bridging the α-and β-phosphate now interacts with S78, with the β-phosphate facing away from the polyP tunnel rather than towards it and the α-phosphate is positioned halfway between the two AMP confomers (Figure 7F, superimposition of all substrates Figure 8B).

We also determined a crystal structure of *Lf*PPK2 in complex with TMP, a 2′-deoxy pyrimidine NMP. The nucleobase shows two ionic interactions with E117 *via* the NH group in position 3 and with the oxygen in the C4-position (Figure 8A). E117 is similarly identified as the base-binding residue in the AMP/ADP-bound structures. The position of the ribofuranose ring strongly deviates from the one in the purine nucleotide-bound structures (Figure 8B), with a distance of 4.2 Å between the hydroxy group of TMP and the C3′ hydroxy group of AMP. This position places the ring too far away from other residues for further interactions. The phosphate group of TMP interacts with R108 at a distance of 2.4 Å. In the AMP and ADP-bound structures, this residue interacted with the C2′ hydroxyl group of the ribose which is not present in TMP. The phosphate of TMP is oriented towards the polyP tunnel. TMP may bind less tightly to the active site, due to the lack of interaction with the ribofuranose ring, which could in part explain the observed lower level of phosphorylation of TMP in comparison to purine nucleotides. Since we did not obtain structures with pyrimidine ribonucleotides (e.g. CMP, UMP), we cannot compare the binding of the ribose rings at this point.

The structures reported here provide novel insights into the substrate binding mode of PPK2-II enzymes and are the first solved substrate-bound structures of PPK2 enzymes from subclass II as well as the first with a 2′-deoxy nucleotide or a pyrimidine-derived nucleotide of any PPK2 enzyme.

## Conclusion

*Bc*PPK2 and *Lf*PPK2 have an exceptionally broad substrate range, encompassing purine, pyrimidine, and 2’-deoxy NMPs. This substrate flexibility not only underlines their potential for various applications in nucleotide synthesis but also positions them as valuable tools in biotechnological and therapeutic research. Their ability to efficiently use different nucleotide substrates makes them particularly promising for developing novel nucleoside analogues and improving metabolic pathways. A recent structure-guided rational design campaign on a PPK2-III enzyme shows that exchange of single amino acids close to the active site can have a great impact on the substrate promiscuity towards non-canonical nucleotides (shift from adenosine-derived to guanosine-derived nucleotides).^[39]^ Consequently, the structures solved can act as a starting point for further structure-guided tailoring of PPK2-II enzymes.

Our results further highlight that the PPK2-II’s selectivity to catalyse the formation of NDPs from NMPs is not only a matter of enzyme kinetics but strongly dependent on the properties of the starting substrate. Both enzymes catalyse NTP formation exclusively with purine nucleotides. The successful application of the enzymes for preparative synthesis of the pyrimidine nucleotide CDP was demonstrated. Taken together, these results provide a basis for future studies in order to further tailor easy-to-adapt PPK2 biocatalysts to specific needs including potential applications in the synthesis of pharmaceutical compounds.

In summary, this work extends the enzymatic toolbox available for synthesis of (non-)canonical nucleotides in various phosphorylation states, with PPK2-II enzymes as efficient biocatalysts for NDP synthesis.

## Experimental Section

### Chemicals

The following chemicals were used: polyP from Merck (average chain length of 25 phosphates), AMP (disodium salt, Sigma Aldrich), GMP (hydrate, TCI), IMP (disodium salt, Sigma Aldrich), XMP (disodium salt, ChemPUR), CMP (disodium salt, Sigma Aldrich), ADP (sodium salt, Sigma Aldrich), GDP (disodium salt, Sigma Aldrich), IDP (trisodium salt, ABCR), CDP (disodium salt, BLD Pharmatech). For cell cultivation, appropriate purity chemicals from Carl Roth (Germany) were used.

### Cloning, Expression and Purification of PPK2-II enzymes

The protein sequences of the PPK2 enzymes were obtained from NCBI. The protein sequences were back translated to codon optimised genes (Invitrogen, Thermofisher) for expression in *E. coli* cells and cloned into a pET28a(+) vector. The gene strings were inserted into linearised vectors using the In-fusion kit (Takara Bioscience) according to the manufacturers protocol. Successful cloning was confirmed by sequencing (Eurofins, Germany). BL21Gold-DE3 cells were transformed with the plasmid and grown on LB-Agar plates containing the appropriate antibiotic. For expression, LB broth was inoculated with an overnight culture of the transformed *E. coli* cells and incubated at 37 °C while shaking at 160 rpm. Optical density was measured frequently, at values of OD_600_∼0.5 protein production was induced by adding IPTG (final concentration of 0.1 mM). Cells were further cultured at 20 °C for 20 hours then harvested by centrifugation. To isolate the overproduced enzymes, cells were lysed by sonication followed by centrifugation to remove cell debris and unlysed cells. Thereafter, the cleared lysate was subjected to Ni-NTA affinity chromatography. To remove imidazole from the preparation, the purified enzyme was subjected to size exclusion chromatography using a prepacked PD10 column (GE Healthcare). For crystallisation experiments, the enzyme preparation was further purified by semi preparative size exclusion using a Sephadex S200 column.

### Enzymatic Assays

polyP dependent phosphorylation of nucleotides experiments contained the following: 50 mM Tris-HCl (pH 8.0), 10 mM MgCl2, 10 mM polyP (calculated as single phosphates), 1 mM nucleotide and 1 µM enzyme in a 1 mL reaction scale. The reactions were prepared on ice, started by addition of the enzyme and incubated at 37 °C while shaking at 350 rpm in a thermo cycler. Samples were quenched *via* addition of HClO_4_ (final concentration 2.5 %), frozen in liquid nitrogen and stored at -20 °C until analysis. All assays were performed at least in triplicates with standard deviations indicated by error bars in the corresponding figures.

### HPLC-Analysis

HPLC analysis was carried out on an Agilent 1260 Infinity II system equipped with an ISAsphere 100-5 C18 column (250 mm, 4 mm, 5 µm). A gradient method was used with 1 mL/min flowrate and sodium acetate buffer A (40 mM, pH 4.2) and acetonitrile B. The gradient was 10 min 98% A, 6 min 98% → 70% A, 2 min 70% A, 2 min 70% A → 98% A, 10 min 98% A. The detection wavelength was set to 270 nm.

### CE-ESI-MS Analysis

CE-ESI-MS measurements were adapted from previous published works and performed on an Agilent 7100 CE system (Agilent Technologies) coupled to an Agilent G6495C Agilent QQQ mass spectrometer equipped with a commercial Agilent jet stream (AJS) electrospray ionization (ESI) source and an Agilent liquid coaxial interface.^[34,35]^ To achieve a stable ESI spay, sheath liquid containing a _dd_H_2_O-isopropanol mixture (1:1 v-%) was pumped with an Agilent 1200 isocratic LC pump and a 1:100 splitter, which resulted in a flow of 10 µl·min^-1^. Separation was performed on a bare fused silicia capillary (100 cm length, 50 μm internal diameter) activated and flushed with 1 M NaOH and water for 10 minutes before the first measurement, respectively. Before each measurement, the capillary was washed with _dd_H_2_O (200 s) and background electrolyte of 35 mM ammonium acetate, titrated to pH 9.75 with NH_4_OH (300 s). Samples were applied via a pressure of 100 mbar for 10 s. Separation was achieved with the background electrolyte and a CE separation voltage of +30 kV, resulting in a stable current of 22 µA. MS were performed in negative ionization mode and the following parameters: capillary voltage: -2000 V; nebulizer gas: 8 psi; gas temperature: 150 °C and a flow of 11 L/min; sheath gas temperature: 175 °C and a flow of 8 L/min. Nozzle voltage was set to 2000 V and pressure RF was between 60 V and 90 V. Peak assignment was done via MS/MS transitions (MS parameters in Table S3).

### NDP preparation

Enzymatic reactions used for nucleotide purification contained 50 mM Tris-HCl (pH 8.0), 10 mM MgCl2, 10 mM polyP (calculated as single phosphates), 10 mM CMP and 5 µM PPK2 enzyme in a 5 mL reaction scale. The reactions were prepared on ice, started by addition of the enzyme and incubated at 37 °C while shaking at 350 rpm in a thermoshaker. Reactions were stopped by heating to 85 °C for 5 minutes. Afterwards, the sample was centrifuged (5400 g, 4 °C, 10 min) and applied onto a HiPrep Q FF 16/10 column (20 mL, Cytiva) connected to an NGC Chromatography System (Bio-Rad). The column was preequilibrated with _dd_H_2_O. Nucleotides were eluted with a gradient of 500 mM NaClO_4_ (pH 4.2). Fractions containing the desired NDP were pooled and lyophillised until dry. Afterwards the sample was dissolved in 1 mL 500 mM NaClO_4_ and added to 49 mL chilled aceton (− 20 °C) and stored at − 20 °C for 1.5 h to allow precipitation of the NDP salt. Next, the sample was centrifuged (5400 g, 4 °C, 30 min), the precipitate washed with chilled aceton (40 mL) followed by a additional centrifugation step (15 min). The supernatant was discarded and the product was dried at room temperature to allow evaporation of residual acetone. The amount of NDP in the final product was determined via UV/Vis spectroscopy (Jasco V-730). Concentrations were calculated according to the Beer-Lambert Law using an molar extinction coefficient of 8.9 mM^−1^ · cm^−1^ for CDP.

### Crystallisation and Soaking

Crystallisation assays leading to structures with the PDB IDs 9H8J, 9H8K and 9H8L were set up using an Oryx Nano liquid dispenser (Douglas Instruments, East Garston, UK) using sitting drop vapour diffusion in MRC 2-Well UVP plates, SWISSCI, Buckinghamshire, UK) at 20 °C. Crystals of apo *Lf*PPK2 formed after 5 days in wells containing 0.30 µL of protein solution (6.0 mg/mL) and 0.30 µL of a reservoir solution containing 15% (w/v) PEG 3,350 and 0.1 M succinic acid buffer at pH 7.0. These crystals were cryoprotected with a mixture of reservoir solution and 10% (v/v) 2*R*,3*R*-(–)-butanediol, mounted on a nylon loop, and flash-cooled in liquid nitrogen. Additional crystals of apo *Lf*PPK2, suitable for soaking experiments, formed after 7–10 days in wells containing 0.25 µL of protein solution (6.0 mg/mL) and 0.40 µL of reservoir solution with 1.5 M ammonium sulfate and 0.1 M sodium acetate buffer (pH 4.6). The *Lf*PPK2-AMP and *Lf*PPK2-TMP complexes were prepared by soaking apo *Lf*PPK2 crystals in solutions containing AMP and TMP, respectively. For AMP soaking, the reservoir solution was mixed with 10% (v/v) of AMP solution (100 mM final concentration, in 40 mM Tris/HCl buffer, pH 8.0). For TMP soaking, the reservoir solution was mixed with 15% (v/v) TMP solution (750 mM final concentration, in 40 mM Tris/HCl buffer, pH 8.0). The apo *Lf*PPK2 crystals were soaked in these solutions for 24 h, cryoprotected with a mixture of reservoir solution and 15% (v/v) 2*R*,3*R*-(–)-butanediol, mounted on a nylon loop, and flash-cooled in liquid nitrogen. Enzyme crystals leading to structures with the PDB IDs 9GIA and 9GP9 were prepared at a concentration of 10 mg·mL^−1^ using the sittingdrop vapour diffusion method at room temperature. *Lf*PPK2-II was crystallised in a solution containing 100 mM MES/NaOH buffer at pH 6.5 and ammonium sulfate at concentrations ranging from 1.2 M to 1.5 M. *Bc*PPK2-II crystals were grown in the presence of 100 mM Tris/HCl at pH 8.5, 200 mM lithium sulfate, and 22% (w/v) PEG 4000. Apo-enzyme crystals suitable for diffraction were transferred into reservoir liquor containing 25% PEG 4000 for *Bc*PPK2-II crystals, while 4 M ammonium sulfate was used as a cryoprotectant for *Lf*PPK2-II crystals. All crystals were flash-frozen in liquid nitrogen. For soaking experiments, nucleotide solutions were added to the mother liquor in a 1:1 (v/v) ratio to reach a final concentration of 10 mM, and crystals were flash-frozen similarly.

### Data Collection and Structure Determination

X-ray diffraction data for the *Lf*PPK2 apo structure (PDB 9H8J) and the *Lf*PPK2-AMP complex (PDB 9H8K) were collected on the ID30A-3 (MASSIF-3) beamline at the European Synchrotron Radiation Facility (ESRF, Grenoble, France) using an EIGER X 4M detector. Data for the *Lf*PPK2-TMP complex (PDB 9H8L) were collected on the ID30B beamline at the ESRF with an EIGER2 X 9M detector. Diffraction data for the *Bc*PPK2 apo structure (PDB 9GIA) and *Lf*PPK2-ADP complex structure (PDB 9GP9) were collected at the PXI-X06SA beamline at the Paul Scherrer Institute (PSI, Villigen, Switzerland) using a DECTRIS EIGER2 X 16M detector and maintained at 100 K with a nitrogen stream. The datasets were processed using autoPROC^[41]^and scaled with Aimless^[42]^. Molecular replacement was conducted with Phaser^[43]^, employing either the Alphafold model of *Lf*PPK2 (Uniprot A0A1E4R1F9) or the *Pa*PPK2-II structure (PDB 3CZP) as the search model for PDB entrys 9H8J, 9H8K, 9H8L or 9GIA, 9GP9 respectively. Models were built and refined iteratively using COOT^[44]^ and either REFMAC^[45]^ or Phenix.refine.^[46]^ Restraints for AMP and TMP were generated with the Grade Web Server (Global Phasing Ltd., UK). The electron density for TMP in the *Lf*PPK2-TMP (PDB 9H8L) structure was well defined, while weak electron density for AMP in the catalytic site of the *Lf*PPK2-AMP (PDB 9H8K) structure indicated partial occupancy, with two distinct AMP conformations observed. Final structure validation was performed using MolProbity.^[47]^ Data collection and refinement statistics are provided in Table S1 and Table S2. Figures were prepared using PyMOL (The PyMOL Molecular Graphics System, Version 2.6 Schrödinger, LLC).

## Acknowledgements

This work was supported by the Deutsche Forschungsgesellschaft (DFG, German Research Foundation, Heisenberg programme grant no. 527572100 to JNA) and CIBSS (EXC-2189, Project ID 390939984 to HJJ). We thank all members of the European Synchrotron Radiation Facility and the beamline at the Paul Scherrer Institute for support in crystallographic data collection. We thank Dr. Philippe Bisel and Sascha Ferlaino) for conducting NMR measurements. Dr. Wolfgang Hüttel is thanked for his support with bioinformatics. Chiara Schmid is acknowleged for her initial work with the PPK2 enzymes during her bachelor thesis. Emely Jockmann is thanked for proof-reading and commenting on the draft of this manuscript.

## Notes

### Competing Interest Statement

The authors have declared no competing interest.

