## Supplemental Information for "Structural Insights into Broad-Range Polyphosphate Kinase 2-II Enzymes Applicable for Pyrimidine Nucleoside Diphosphate Synthesis"

**for**

### Supplemental Tables

**Table S 1.** Crystallographic data collection and refinement statistics of structures with the PDB IDs 9GIA and 9GP9.

| Structure | *Bc*PPK2-II apo | *Lf*PPK2-II ADP |
|---|---|---|
| PDB entry | 9GIA | 9GP9 |
| **Data Collection** | | |
| Wavelength (Å) | 1.00 | 1.00 |
| Space group | *I* 2 2 2 | *I* $4_1$ 2 2 |
| *Cell Dimensions* | | |
| a, b, c (Å) | 93.13, 116.82, 130.54 | 109.78,109.78, 286.68 |
| α, β, γ (Å) | 90, 90, 90 | 90, 90, 90 |
| Resolution (Å) | 48.6 - 2.55 | 48.39 - 2.88 |
| High res. (Å) | 2.67 - 2.55 | 3.03 - 2.88 |
| $R_{merge}$ | 0.12 (1.07) | 0.09 (1.54) |
| $R_{pim}$ | 0.07 (0.84) | 0.04 (0.78) |
| $I / \sigma_I$ | 8.6 (1.0) | 16.4 (1.5) |
| Completeness (%) | 99.67 (99.52) | 98.58 (98.49) |
| Multiplicity | 6.4 (4.4) | 8.9 (9.1) |
| $CC_{1/2}$ | 0.998 (0.53) | 0.99 (0.58) |
| Wilson B-factor | 64.49 | 99.13 |
| **Refinement** | | |
| No. of unique reflections | 23,521 (2907) | 20,084 (2802) |
| Reflections used in $R_{free}$ | 1,177 (145) | 1,004 (140) |
| $R_{work}/R_{free}$ | 0.21/0.25 | 0.20/0.23 |
| *Number of atoms* | | |
| macromolecules | 4288 | 3341 |
| ligands | 48 | 84 |
| solvent | 56 | 4 |
| *R.m.s deviations* | | |
| Bond lengths (Å) | 0.010 | 0.012 |
| Bond angles (°) | 1.72 | 2.00 |
| *B-factors* | | |
| macromolecules | 63.57 | 92.44 |
| ligands | 97.40 | 130.83 |
| solvent | 52.58 | 73.87 |
| *Ramachandran - Φ, Ψ angle distribution for residues (%)* | | |
| favoured | 97.59 | 97.21 |
| allowed | 1.81 | 2.79 |
| outliers | 0.60 | 0.00 |

**Table S 2.** Crystallographic data collection and refinement statistics of structures with the PDB IDs 9H8J, 9H8K, 9H8L.

| Structure | ***Lf*PPK2 apo** | ***Lf*PPK2-AMP** | ***Lf*PPK2-TMP** |
|---|---|---|---|
| PDB entry | 9H8J | 9H8K | 9H8L |
| **Data Collection** | | | |
| Space group | $P\ 2_1$ | $P\ 6_222$ | $P\ 6_222$ |
| *a, b, c* (Å) | 86.11, 71.93, 97.69 | 160.77, 160.77, 67.72 | 159.60, 159.60, 67.43 |
| *α, β, γ* (deg) | 90.00, 92.59, 90.00 | 90.00, 90.00, 120.00 | 90.00, 90.00, 120.00 |
| Wavelength (Å) | 0.9677 | 0.9677 | 0.8731 |
| Resolution (Å) | 66.03 – 2.20 (2.26 – 2.20) | 69.62 – 2.10 (2.16 – 2.10) | 138.21 – 2.10 (2.16 – 2.10) |
| Molecules per asymmetric unit | 4 | 1 | 1 |
| Total/unique no. of reflections | 413,086 (59,963) | 1,171,782 (30,456) | 1,138,655 (29,017) |
| $R_{merge}{}^{a,b}$ | 0.199 (1.255) | 0.243 (3.778) | 0.123 (2.471) |
| $R_{pim}{}^{a,c}$ | 0.080 (0.510) | 0.040 (0.599) | 0.020 (0.385) |
| $CC_{1/2}{}^{a,d}$ | 0.992 (0.632) | 0.999 (0.629) | 0.999 (0.763) |
| $I/\sigma(I)\ ^a$ | 8.6 (1.4) | 17.7 (1.7) | 24.8 (2.4) |
| Redundancy[a] | 6.9 (7.0) | 38.5 (40.6) | 39.2 (41.9) |
| Completeness (%)[a] | 98.7 (99.2) | 99.5 (94.8) | 96.6 (100.0) |
| **Refinement** | | | |
| No. of reflections used in refinement/test set | 59,934 (5,993) | 30,398 (2,865) | 28,804 (2,919) |
| $R_{work}{}^e$ | 0.204 (0.260) | 0.193 (0.263) | 0.193 (0.267) |
| $R_{free}{}^f$ | 0.253 (0.312) | 0.217 (0.308) | 0.208 (0.326) |
| Number of Atoms[g] | 8601 | 2321 | 2156 |
| protein | 8087 | 2108 | 2032 |
| ligands | 48 | 86 | 36 |
| solvent | 466 | 127 | 88 |
| Average *B*-Factors (Å$^2$) | 41.8 | 47.1 | 50.5 |
| protein | 42.0 | 46.4 | 50.3 |
| ligands | 42.1 | 65.2 | 66.9 |
| solvent | 37.3 | 47.0 | 48.8 |
| RMS Deviations | | | |
| bonds (Å) | 0.0059 | 0.0076 | 0.0060 |
| angles (deg) | 0.90 | 0.79 | 0.71 |
| Ramachandran plot (%)[h] | | | |
| favoured | 97.16 | 97.17 | 97.47 |
| allowed | 2.84 | 2.83 | 2.53 |
| outliers | 0 | 0 | 0 |

[a] Values in parentheses refer to the highest-resolution shell of the data. [b] $R_{merge} = \sum|I_h - \langle I_h\rangle|/\sum\langle I_h\rangle$; $I_h$ = intensity measure for reflection *h;* $\langle I_h\rangle$ = average intensity for reflection *h* calculated from replicate data. [c] $R_{pim} = \sum(1/(n-1))^{1/2}|I_h - \langle I_h\rangle|/\sum\langle I_h\rangle$; *n* = number of observations (redundancy). [d] $CC_{1/2} = \sigma_\tau^2/(\sigma_\tau^2 + \sigma_\varepsilon^2)$, where $\sigma_\tau^2$ is the true measurement error variance and $\sigma_\varepsilon^2$ is the independent measurement error variance. [e] $R_{work} = \sum||F_o| - |F_c||/\sum|F_o|$ for reflections contained in the working set. $|F_o|$ and $|F_c|$ are the observed and calculated structure factor amplitudes, respectively. [f] $R_{free} = \sum||F_o| - |F_c||/\sum|F_o|$ for reflections contained in the test set held aside during refinement. [g] Per asymmetric unit. [h] Assessed by MolProbity.

**Table S 3.** MS parameter used for detection of AMD, ADP and ATP.

| Compound | MRM transition | Type of transition | Collision Energy [V] | Cell Acceleration [V] |
|---|---|---|---|---|
| ATP | 506.0 → 407.9 | $[M\text{-}H]^- \rightarrow [M\text{-}H\text{-}H_2PO_4]^-$ | 25 | 1 |
| | 506.0 → 158.9 | $[M\text{-}H]^- \rightarrow [P_2O_6H]^-$ | 37 | 5 |
| ADP | 426.0 → 328.0 | $[M\text{-}H]^- \rightarrow [M\text{-}H\text{-}H_3PO_4]^-$ | 17 | 4 |
| | 426.0 → 79.0 | $[M\text{-}H]^- \rightarrow [PO_3]^-$ | 54 | 4 |
| AMP | 346.1 → 134.1 | $[M\text{-}H]^- \rightarrow [C_5H_4N_5]^-$ | 41 | 4 |
| | 346.1 → 78.9 | $[M\text{-}H]^- \rightarrow [PO_3]^-$ | 46 | 4 |

### Supplemental Figures

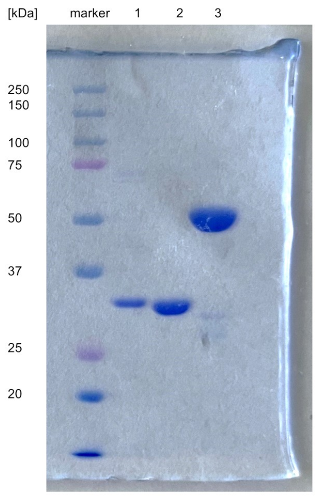

**Figure S1.** SDS gel of purified enzymes. Lane 1: *Bc*PPK2 (34.4 kDa), lane 2: *Lf*PPK2 (33.9 kDa), AjPPK2 (58.0 kDa)

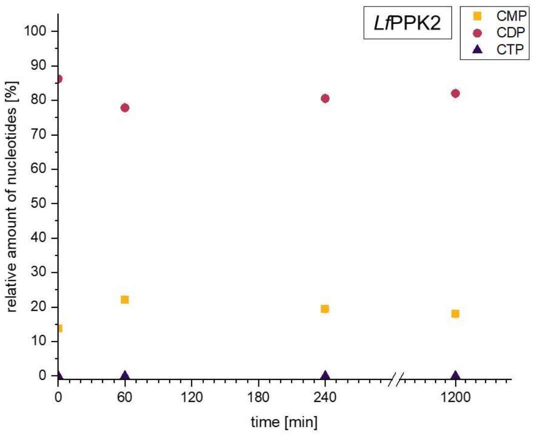

**Figure S2.** Time course of CDP consumption catalysed by *Lf*PPK2. Assays comprised of 1 mM CDP, 1 µM *Lf*PPK2, 10 mM polyP, 10 mM $MgCl_2$. in 50 mM Tris buffer (pH 8.0). Samples were taken after 0, 10, 30, 60, 120, 240 and 1200 minutes and subjected to HPLC analysis.

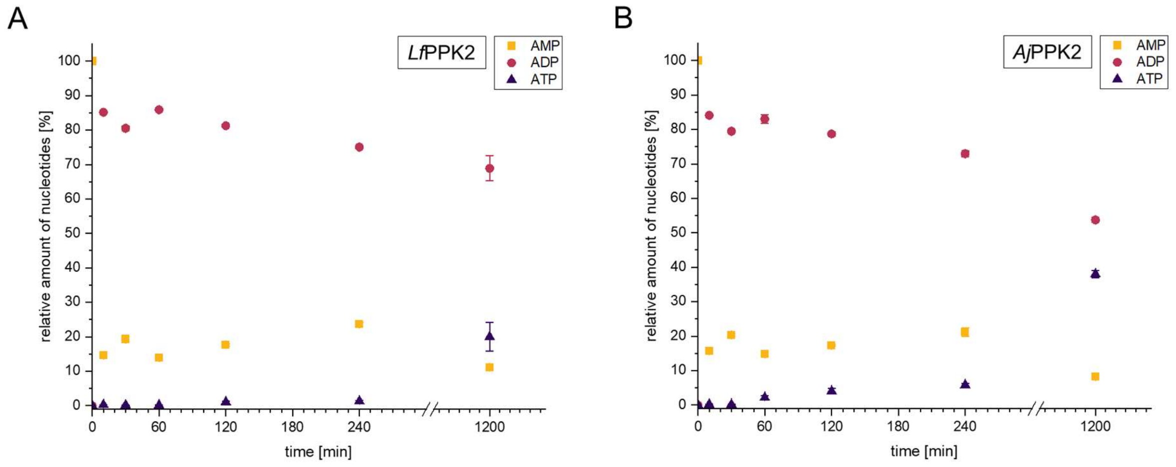

**Figure S3.** Time course of ADP and ATP formation catalysed by *Lf*PPK2 (A) and *Aj*PPK2 (B). Assays comprised of 1 mM AMP, 1 µM PPK2 enzyme, 10 mM polyP, 10 mM $MgCl_2$. Samples were taken after 0, 10, 30, 60, 120, 240 and 1200 minutes and subjected to HPLC analysis.

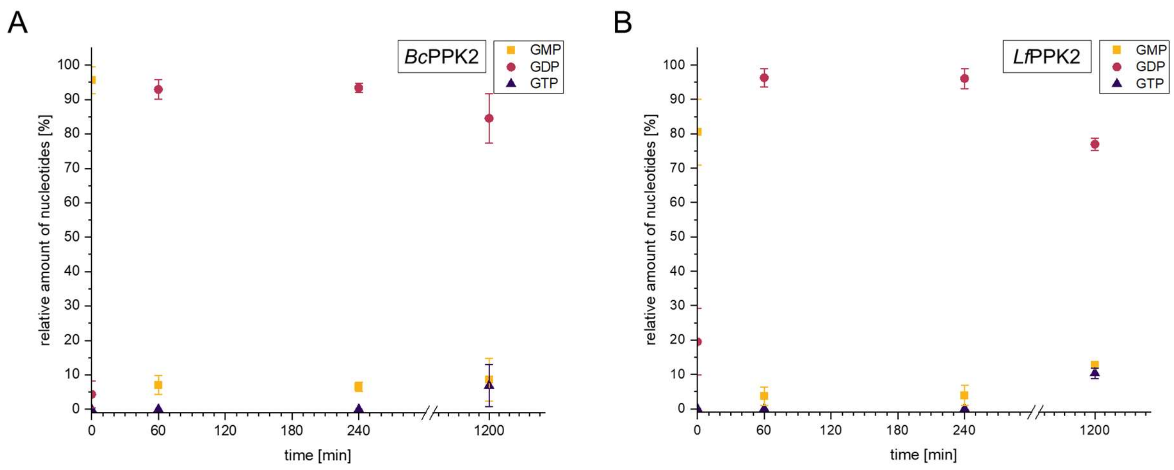

**Figure S4.** Time course of GDP and GTP formation catalysed by *Bc*PPK2 (A) and LfPPK2 (B). Assays comprised of 1 mM GMP, 1 µM PPK2 enzyme, 10 mM polyP, 10 mM $MgCl_2$. Samples were taken after 0, 60, 240 and 1200 minutes and subjected to HPLC analysis.

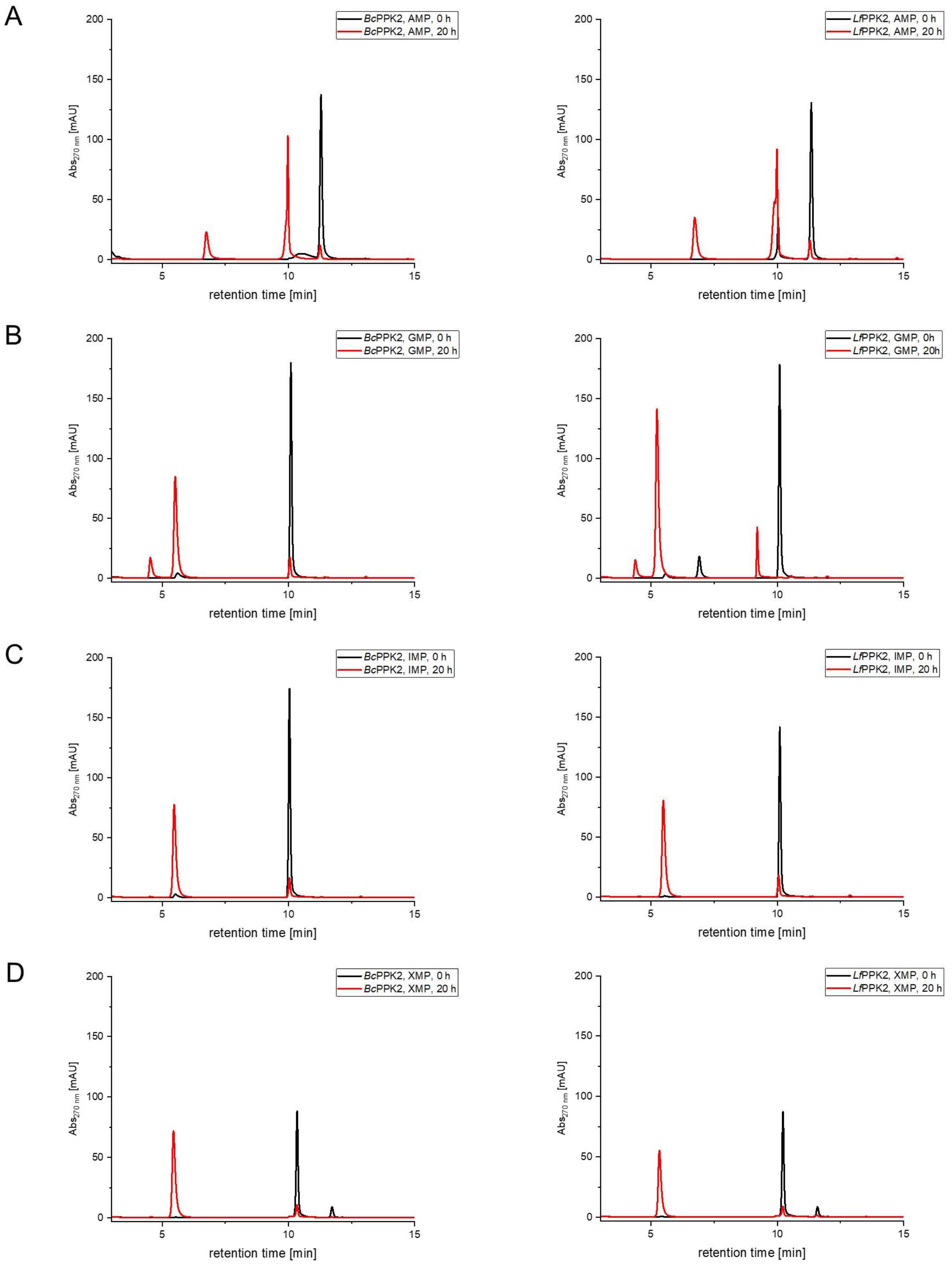
A
BcPPK2, AMP, 0 h
BcPPK2, AMP, 20 h
LfPPK2, AMP, 0 h
LfPPK2, AMP, 20 h
B
BcPPK2, GMP, 0 h
BcPPK2, GMP, 20 h
LfPPK2, GMP, 0h
LfPPK2, GMP, 20h
C
BcPPK2, IMP, 0 h
BcPPK2, IMP, 20 h
LfPPK2, IMP, 0 h
LfPPK2, IMP, 20 h
D
BcPPK2, XMP, 0 h
BcPPK2, XMP, 20 h
LfPPK2, XMP, 0 h
LfPPK2, XMP, 20 h
Abs270 nm [mAU]
retention time [min]
0
50
100
150
200
5
10
15

E

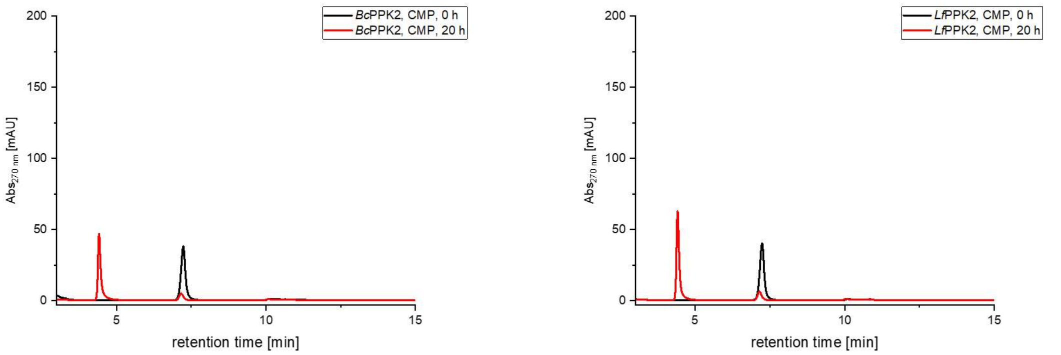

*Bc*PPK2, CMP, 0 h
*Bc*PPK2, CMP, 20 h
*Lf*PPK2, CMP, 0 h
*Lf*PPK2, CMP, 20 h
$Abs_{270\ nm}$ [mAU]
retention time [min]

F

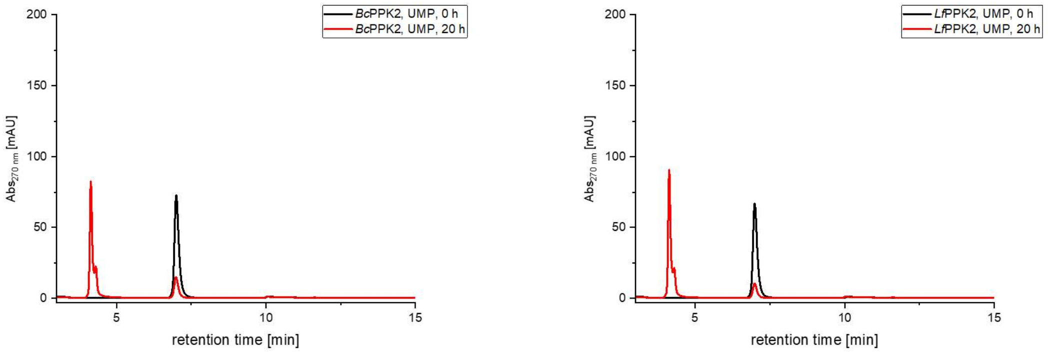

*Bc*PPK2, UMP, 0 h
*Bc*PPK2, UMP, 20 h
*Lf*PPK2, UMP, 0 h
*Lf*PPK2, UMP, 20 h
$Abs_{270\ nm}$ [mAU]
retention time [min]

G

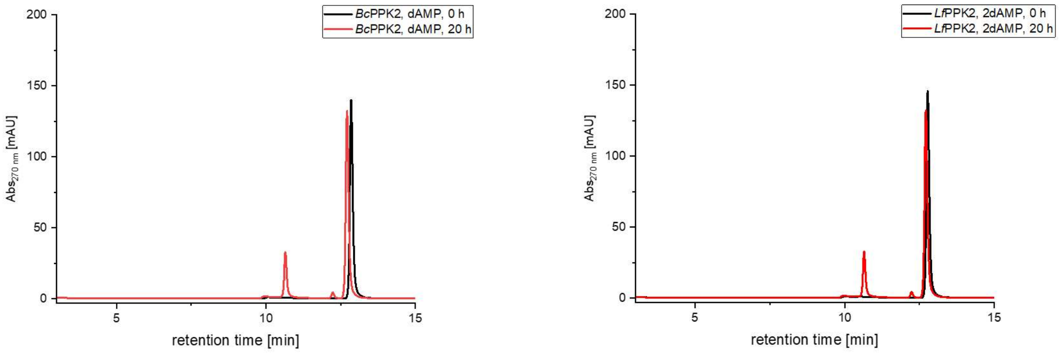

*Bc*PPK2, dAMP, 0 h
*Bc*PPK2, dAMP, 20 h
*Lf*PPK2, 2dAMP, 0 h
*Lf*PPK2, 2dAMP, 20 h
$Abs_{270\ nm}$ [mAU]
retention time [min]

H

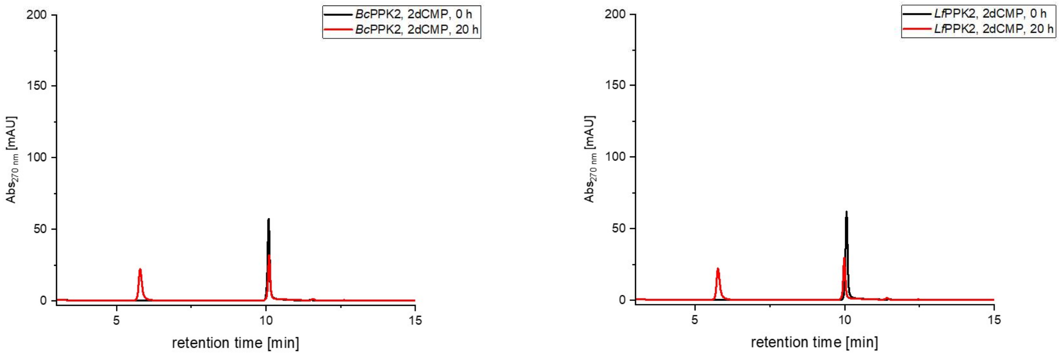

*Bc*PPK2, 2dCMP, 0 h
*Bc*PPK2, 2dCMP, 20 h
*Lf*PPK2, 2dCMP, 0 h
*Lf*PPK2, 2dCMP, 20 h
$Abs_{270\ nm}$ [mAU]
retention time [min]

I

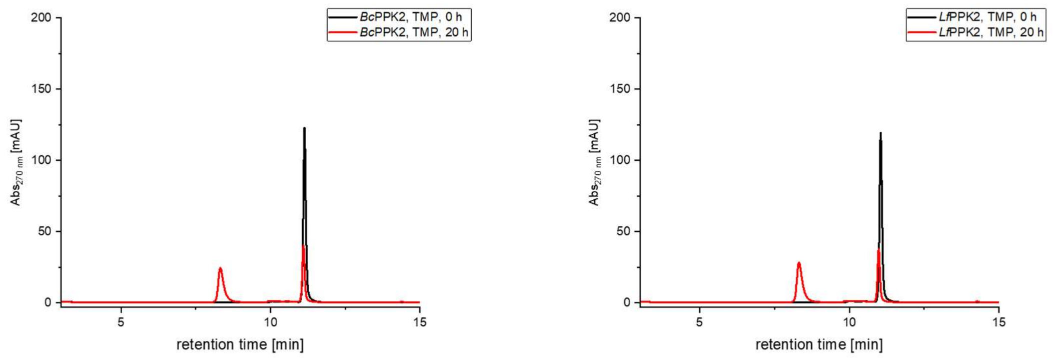

**Figure S5.** HPLC-UV chromatograms of the substrate specifity screening assay for *Bc*PPK2 (left) and *Lf*PPK2 (right). Shown is the 0 h sample (black) and the corresponding sample after 20 h reaction time (red). The following substrates were ised: AMP (A), GMP (B), IMP (C), XMP (D), CMP(E), UMP (F), dAMP (G), dCMP (H), TMP (I). Enzyme assays comprised of 1 µM PPK2 enzyme, 1 mM NMP, 10 mM $MgCl_2$, 10 mM polyP (calculated as single phosphate units) and were incubated at 37 °C for 20 h. After quenching with perchloric acid (2.5% final concentration), samples were subjected to HPLC analysis.

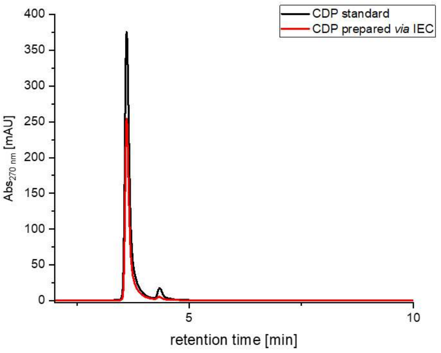

**Figure S6.** HPLC-UV chromatogram of commercial CDP (black, BLD pharm) and *via* ionic exchange chromatography prepared CDP from enzymatic assay (red).

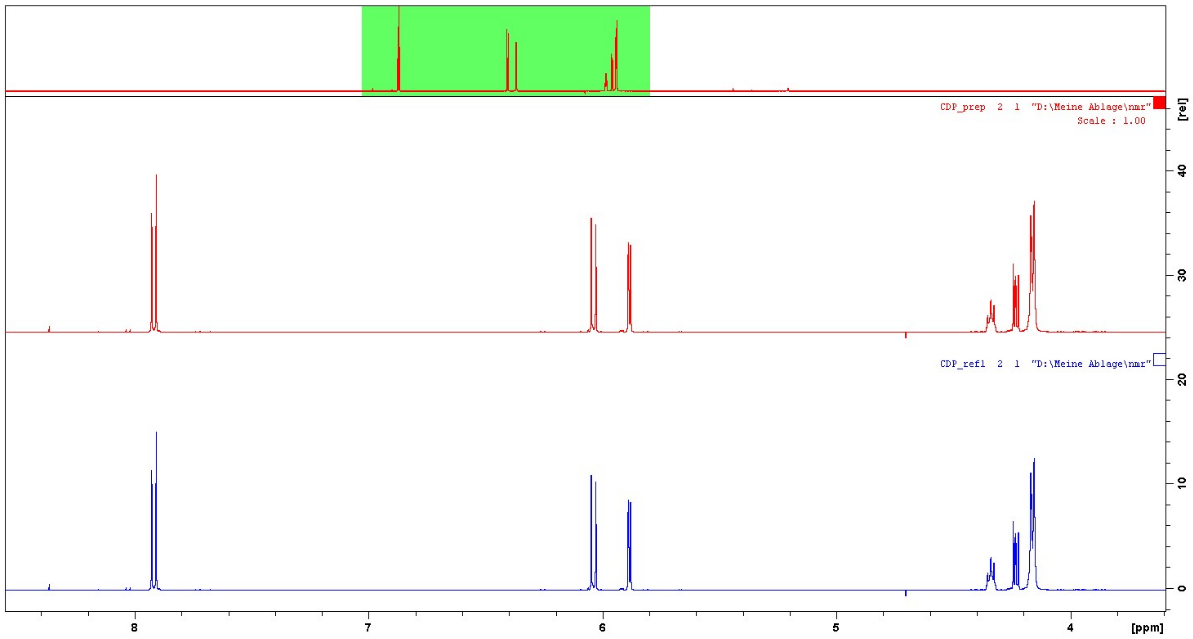

**Figure S 7.** $^1$H NMR spectra comparing CDP reference material (lower spectrum, BLD pharm BD21927) with purified synthesis product (upper spectrum).

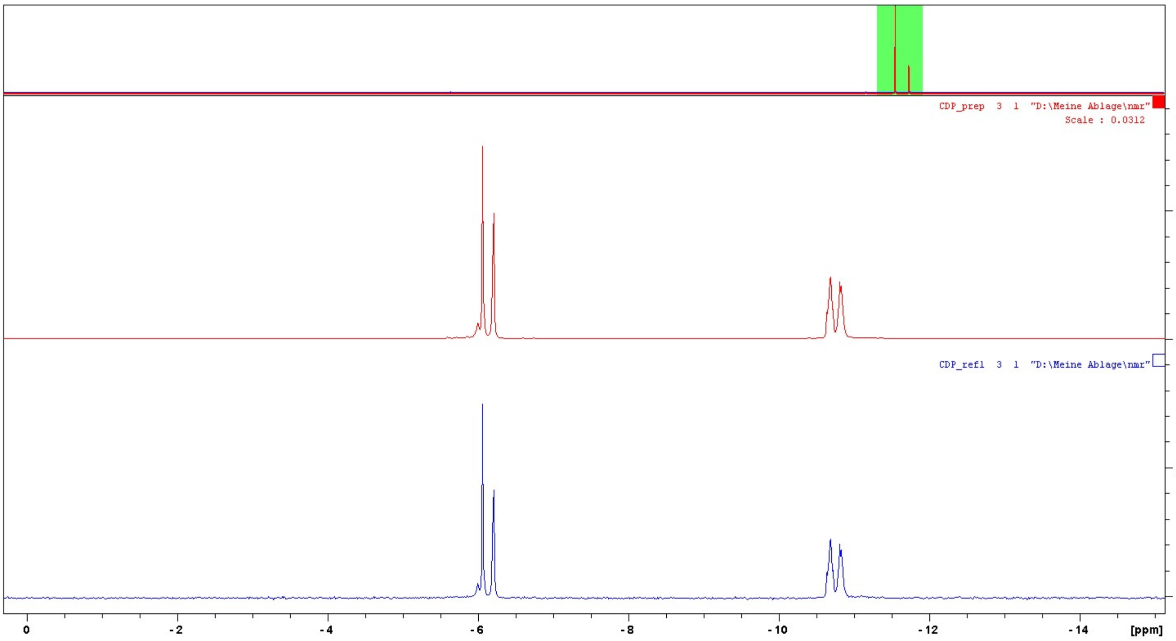

**Figure S 8.** $^{31}P$ NMR spectra comparing CDP reference material (lower spectrum, BLD pharm BD21927) with purified synthesis product (upper spectrum).

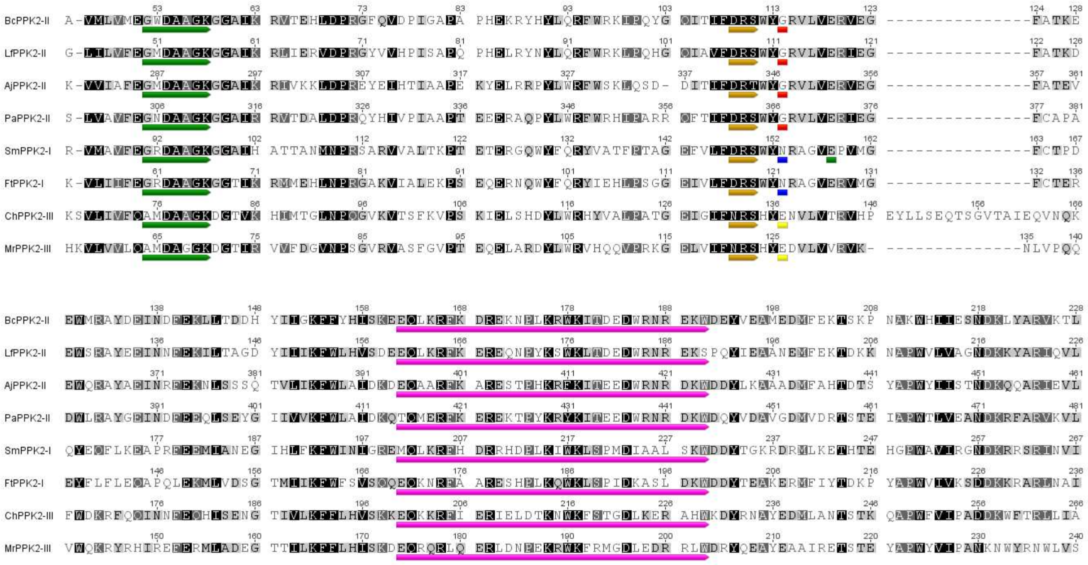

**Figure S 9.** Multiple sequence alignment of PPK2 enzymes of all three subclasses. *Bc*PPK2-II, *Lf*PPK2-II, *Aj*PPK2-II, *Pa*PPK2-II (*Pseudomonas aeruginosa*, UniProt Q9HYF1), *Sm*PPK2-I (*Rhizobium meliloti*, earlier *Sinorhizobium meliloti*, UniProt Q92SA6), *Ft*PPK2-I (*Francisella tularensis*, UniProt Q5NEQ5), *Ch*PPK2-III (*Cytophaga hutchinsonii*, UniProt A0A6N4SMB5), *Mr*PPK2-III (*Meiothermus ruber*, UniPRot M9XB82). Characterisic motifs are coloured as follows: green: Walker A motif, gold: Walker B motif, pink: lid domain, red: PPK2-II Gly signature residue, blue: PPK2-I Asn signature residue, yellow: PPK2-III Glu signature residue. Sequence alignment generated with Geneious version 7.1 created by Biomatters.

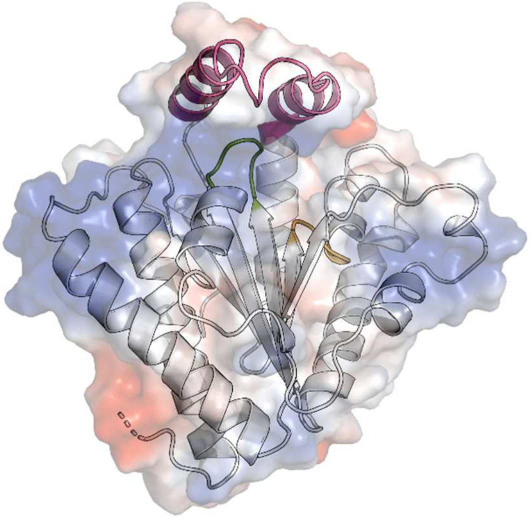

**Figure S 10.** Structural representation of *Lf*PPK2 apo enzyme (PDB ID 9H8J). The electrostatic surface is represented with positively charged residues in blue and negatively charged residues in red. Tertiary structure elements are overlaid in a cartoon representation. Conserved motifs are colour-coded as follows: Walker A, green. Walker B, orange. Lid domain, violet.

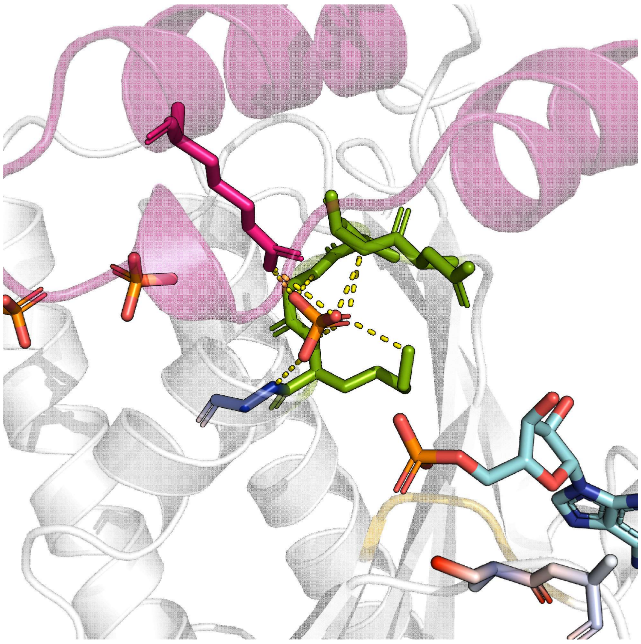

**Figure S 11.** Structural representation of *Lf*PPK2 in complex with AMP (PDB ID 9H8K). A single phosphate unit, located in the polyP tunnel, is coordinated by residues of the Walker A motif (green), as well as the adjacent G57 (blue) and R168 of the lid domain (violet).

#### Enzyme sequences

>*Aj*PPK2

MGSSHHHHHHSSGLVPRGSHMMDTETIASAVLNEEQLSLDLIEAQYALMNTRDQSNAKSLVILVSGIE
LAGKGEAVKQLREWVDPRFLYVKADPPHLFNLKQPFWQPYTRFVPAEGQIMVWFGNWYGDLLATA
MHASKPLDDTLFDEYVSNMRAFEQDLKNNNVDVLKVWFDLSWKSLQKRLDDMDPSEVHWHKLHGL
DWRNKKQYDTLQKLRTRFTDDWQIIDGEDEDLRNHNFAQAILTALRHCPEHEKKAALKWQQAPIPDIL
TQFEVPQAEDANYKSELKKLTKQVADAMRCDDRKVVIAFEGMDAAGKGGAIKRIVKKLDPREYEIHTIA
APEKYELRRPYLWRFWSKLQSDDITIFDRTWYGRVLVERVEGFATEVEWQRAYAEINRFEKNLSSSQ
TVLIKFWLAIDKDEQAARFKARESTPHKRFKITEEDWRNRDKWDDYLKAAADMFAHTDTSYAPWYIIS
TNDKQQARIEVLRAILKQLKADRDTD

>*Bc*PPK2

MGSSHHHHHHSSGLVPRGSHMMENGHLAKVDLTKKIESKSKYNKKLEKYQRRLLALQQILKEEKIAV
MLVMEGWDAAGKGGAIKRVTEHLDPRGFQVDPIGAPAPHEKRYHYLQRFWRKIPQYGQITIFDRSWY
GRVLVERVEGFATKEEWMRAYDEINDFEKLLTDDHYIIGKFFYHISKEEQLKRFKDREKNPLKRWKITD
EDWRNREKWDEYVEAMEDMFEKTSKPNAKWHIIESNDKLYARVKTLKIIISFIEDYFLEHGIELPSYYYE
MKEDIEVLQDVGVKE

>*Lf*PPK2

MGSSHHHHHHSSGLVPRGSHMTQNIKNLDLSIELDKKMYKKKLEVLQYEMLNAQQFLLKNKIGLILVF
EGMDAAGKGGAIKRLIERVDPRGYVVHPISAPQPHELRYNYLQRFWRKLPQHGQIAVFDRSWYGRVL
VERIEGFATKDEWSRAYEEINNFEKILTAGDYIIIKFWLHVSDEEQLKRFKEREQNPYKSWKLTDEDWR
NREKSPQYIEAANEMFEKTDKKNAPWVLVAGNDKKYARIQVLQETLAHIEREALKRGLHLTNVLDKAH
LEDAESSSLNILNEKKKACGRTRAPPPPPLRSGC
